# Self-assembled Star-shaped Chiroplasmonic Gold Nanoparticles for Ultrasensitive Chiro-immunosensor of Viruses

**DOI:** 10.1101/162412

**Authors:** Syed Rahin Ahmed, Éva Nagy, Suresh Neethirajan

**Author notes:** Corresponding author (SN).

## Abstract

Near field optics and optical tunneling light-matter interaction in the superstructure of chiral nanostructures and semiconductor quantum dots exhibit strong optical rotation activity that may open a new window for chiral-based bioanalytes detection. Herein we report an ultrasensitive, chiro-immunosensor using superstructure of chiral gold nanohybrids (CAu NPs) and quantum dots (QDs). Self-assembling techniques were employed to create asymmetric plasmonic chiral nanostructures for extending the spectral range of circular dichroism (CD) response for obtaining superior plasmonic resonant coupling with the QDs excitonic state; this may help to achieve lower the limit of detection (LOD) values. As a result, the designed probe exhibited avian influenza A (H5N1) viral concentration at picomolar level, a significant improvement in sensitivity in comparison to a non-assembled CAu NPs based chiroassay. The practicability of the proposed sensing system was successfully demonstrated on several virus cultures including, avian influenza A (H4N6) virus, fowl adenovirus and coronavirus in blood samples. The results of our study highlights that exciton-plasmon interaction changes chirality and the self-assembled nanostructures are an efficient strategy for enhancing the sensitivity of plasmonic nanosensors.

## 1. Introduction

Modern nanotechnology allows us to systematically manipulate the properties of nanoparticles by heterostructuring various nanomaterials either in one nanoscale structure or in alternating nanostructure surface morphologies, which serves as promising candidates for theoretical understanding, and in commercially viable technologies for biological applications. In particular, heterostructuring of semiconductor nanocrystal/plasmonic nanoparticles (PNPs) has garnered a lot of attention due to: (1) modified emission properties; (2) amplified absorption in the presence of plasmon resonance in the PNPs; and (3) shortening of exciton lifetime because of an increased radiation rate and energy transfer to the PNPs. The manipulation of the aforementioned properties of nanomaterials is vitally important and is considered a key issue for the development of technologies in the areas of nano-optics and nanoplasmonics.^1-6^

Recent years has seen a tremendous number of theoretical and experimental investigations of nanoplasmonics at near field optics with only few studies dealing with nano-chiroptics. The field of nano-chiroptics is a new emerging and an exciting frontier. The popularity of nano-chiroptics has exploded because of novel ways of fabrication of engineered metallic nanostructures with tunable surface morphology and their nano-assembly, which offers unprecedented control over electronic and optical properties^7^. Undoubtedly, light-matter interaction in chiral plasmonic metal-semiconductor nanohybrid structures will introduce a new field in the sub-field of physical optics. The most significant advantage of such nanohybrid structures is that they enhance chiroptical response, which could be of great interest in a variety of applications related to chiral biosensing, opening up new areas of research^8^. In comparison to natural chiral molecules, chiral plasmonic nanostructures not only lead to large chiroptical effects, but they also introduce entirely new concepts of superchiral light in technological applications.^9-11^

In the present study, we assumed that nano-assembly of CAu NPs and QDs through immune-reaction could be used for bioanalytical purposes where chiro-optical properties would be modified by a light-matter interaction. Previously, chiro-optical detection techniques were developed based on assembling achiral plasmonic nanoparticles connected by antigen-antibody bridges, and unexpectedly chirality was created due to either the homo-plasmonic nanoparticles or the twisting of the hetero-plasmonic nanoparticles, thus achieving exceptionally low LODs.^12-15^ Several other methods of nano-assembling plasmonic nanoparticles to generate circular dichroism based on DNA origami,^16,17^ silica nanohelices as chiral templates,^18^ polymerase chain reaction,^19^ DNA scaffolds^20^ and specific non-covalent interactions^21^ have been reported. In addition, the broad understanding of plasmonic structures for enhancing the chiroptical response has been studied both theoretically and experimentally.^22^ Herein, our approach is to enhance chiro-optical properties of CAu NPs by nano-assembling with QDs through antibody-antigen interaction.

Nano-assemblies of chiral plasmonic nanoparticles with QDs may play a fundamental role in an entirely new class of nanohybrids with optical resonances in both near field and far-field coupling mechanisms and with enhanced chiroptical effects^7^. Plasmonic nanostructures could play a vital role in achieving an excellent chiro-optical enhancement from metal semiconductor nanohybrids as well as lower LODs for biosensing applications. Particularly, plasmonic morphological alterations of nanostructure surfaces allows the tailoring of surface Plasmon resonances and hence their interaction with incident light. Another important component is the maximizing of overlapping absorption and emission spectra of CAu NPs and QDs, respectively. Considering the above factors, we have developed a self-assembled star-shaped chiro-plasmonic nanostructure using L(+)-ascorbic acid as a key material. We have demonstrated the utility of this nanostructure by using it in a chiro-immunosensor of avian influenza virus A (H5N1), targeting hemagglutinin (HA) and neuraminidase (NA) surface proteins on virus.

We chose influenza virus A (H5N1) detection as a model analyte because of its high pathogenicity in poultry and poultry consumers. The H5N1 flu strain is of great concern globally due to high mortality rate, continuously expanding host range, and the potential emergence of pandemic diseases. Highly pathogenic avian influenza viruses do not readily transfer between humans; however, the disease can be transmitted from infected domestic poultry to humans via close contact interactions (i.e., interaction with infected feces). The effects of H5N1 can also be felt on a global economic level, as pandemic infections cause large economic losses. Therefore, a rapid and sensitive detection tool for the identification of influenza viruses are crucial for early detection and timely responses to overcome the pandemic threat.^23-26^

In North America, fowl adenoviruses (FAdVs) are responsible for several diseases in chickens namely hepatitis, hydropericardium syndrome, respiratory disease and tenosynovitis, causing significant losses to the poultry industry worldwide every year. An early detection system with high sensitivity and specificity is urgently needed to prevent the spread out of disease in poultry sector.^27^ Infectious bronchitis (IB), a coronavirus, is another important disease of chickens caused by IB virus (IBV) which is one of the primary agents of respiratory disease in chickens worldwide. Chickens susceptible to IBV infection have the signs of gasping, coughing, rales, and nasal discharge, huddling together, appearing depressed as well as wet droppings and increased water consumption.^28^

To demonstrate the real world applications of the proposed chiro-optical sensor, avian influenza A (H4N6), fowl adenoviruses-9 (FAdVs-9) strain and infectious bronchitis virus (IBV) were chosen as bioanalyte target candidates in this study.

## 2. Experimental Section

### 2.1 Materials and reagents

Gold (III) chloride trihydrate (HAuCl_4_·3H_2_O), cetyltrimethylammonium bromide (CTAB), 3,3',5,5'-tetramethylbenzidine (TMB), Hydrogen peroxide (H_2_O_2_), Poly-l-lysine (PLL), Nunc-Immuno 96-well plates, CdTe QDs (emission wavelength 710 nm) were purchased from Sigma-Aldrich (St. Louis, MO, USA). The anti-influenza A (H5N1) virus hemagglutinin (HA) antibody [2B7] (ab135382, lot: GR100708-16), recombinant influenza virus A (Avian/Vietnam/1203/04) (H5N1) (lot: GR301823-1), goat anti-mouse IgG, horseradish peroxidase (HRP)-conjugated whole antibody (Ab 97023, lot: GR 250300-11) and immunoassay blocking buffer (Ab 171534, lot: GR 223418-1) were purchased from Abcam, Inc. (Cambridge, UK). Recombinant influenza virus A (H1N1) (California) (CLIHA014-2; lot: 813PH1N1CA) was purchased from Cedarlane (Ontario, Canada). Influenza A (H5N2) hemagglutinin antibodies (Anti-H3N2 antibodies HA MAb, Lot: HB05AP2609), Influenza A (H7N9) hemagglutinin antibodies (Anti-H7N9 antibody HA MAb, Lot: HB05JA1903), recombinant influenza virus A (H5N2) HA1 (A/Ostrich/South Africa/A/109/2006)(lot: LC09AP1021), recombinant influenza virus A (H7N8) HA1 (A/Mallard/Netherlands/33/2006) (lot: LC09AP1323) and recombinant influenza virus A (H7N9) HA1 (A/Shanghai/1/2013) (lot: LC09JA2702) were purchased from Sino Biological, Inc. (Beijing, China). Avian influenza H5N1 neuraminidase polyclonal antibody (Cat. PA5-34949) was received from Invitrogen (Ontario, Canada). Anti-H4 (A/environment/Maryland/1101/06)(H4N6) polyclonal antibody was purchased from MyBioSource Inc., San Diego, USA. Chicken whole blood (Cat. No: IR1-080N) was received from Innovation Research, Michigan, USA. All experiments were performed using highly pure deionized (DI) water (>18 MΩ·cm).

### 2.2 Preparation of different shaped CAu NPs

Chiral gold nanoparticles (CAu NPs) with different shapes were prepared by varying L(+)- ascorbic acid concentrations. 20 μL of varying concentrations of L(+)-ascorbic acids (0.1, 0.05, 0.01, & 0.005 M) was mixed with 1 mL of 20 mM aqueous HAuCl_4_ solution separately under vigorous stirring at 25 °C. Color appeared within a few seconds based on L(+)-ascorbic acids concentration indicating nanoparticle formation. Here, L(+)-ascorbic acids acts as both the reducing agent and as a stabilizer as well as chiral ligands for CAu NPs.

### 2.3 Preparation of self-assembly chiral gold nanostructure

Self-assembled structures of CAu NPs were prepared as previously reported with slight modifications.^12^ For example, 10 mL of 2.5×10^−4^ M HAuCl_4_ and 0.05 M CTAB were mixed and gently stirred for 10 min. Then a 0.01 M (50 μL) solution of L(+)-ascorbic acid was added to the mixture followed by the addition of 20 μL AgNO_3_ (0.01 M) solution. Self-assembled nanostructures were separated from solution by centrifugation at 2000 rpm for 10 min, and were redispersed in 1 mL water.

### 2.4 Avian influenza A H4N6 virus culture

Low pathogenic avian influenza virus A H4N6 (Avian influenza virus A/Duck/Czech/56 (H4N6)) was propagated in 11-day-old embryonated chicken eggs by inoculation into the allantoic cavity.^29^ Virus titer in allantoic fluid was determined at 72 h post-inoculation and expressed as 50% tissue culture infective dose of 128 HAU/50 μL.

### 2.5 Fowl adenoviruses-9 (FAdVs-9) virus culture

FAdV-9 strain was propagated in chicken hepatoma cells (CH-SAH cell line) as described previously.^30^ CH-SAH cells were grown to confluency at 37°C, 5% CO_2_ in Dulbecco’s modified Eagle’s medium/nutrient mixture F-12 Ham with 10% non-heat-inactivated fetal bovine serum as described previously. After infection with the virus, the cells were fed with maintenance medium of D-MEM/F-12 with all the additions, except that the FBS was reduced to 5%. FAdV-9 viral titer in allantoic fluid was determined as 5X10^7^ PFU/mL.

### 2.6 Infectious bronchitis virus (IBV) culture

The viruses were propagated and titrated as previously described.^31,32^ The procedures were carried out in specific pathogen free (SPF) embryonated eggs and the titers were determined by the method of Reed and Muench.^33^ The stock solution of viral titers was 1 ×10^6^ EID_50_/mL.

### 2.7 Specificity of antibodies towards target influenza A (H5N1) virus

Conventional ELISA method was performed to check the specificity of the anti-H5N1 HA (Ab 135382) for influenza virus A/ Vietnam 1203/04/2009 (H3N2). A total of 50 μL (1μg/mL) of virus solution in PBS buffer (pH 7.5) was added to PS plate and kept overnight at 4°C. The next day, wells were rinsed three times with PBS buffer (pH 7.5), immunoassay blocking buffer (Ab 171534, 100 μL) was added and kept at room temperature for 2h. After rinsing three times with PBS buffer (pH 7.5) solution, anti-H5N1 HA Ab (1 μg/mL), anti-H5N1 NA antibody (1 μg/ml), anti-H5N2 HA antibody (1 μg/ml), and anti-H7N9 HA antibody (1 μg/ml) were added to each of the wells, and the plate was incubated at room temperature for 1 h. Again it was washed with PBS buffer (pH 7.5), and HRP-labelled secondary antibody (50 μL, 1 μg/mL) was added and incubated at room temperature for 1h. Then unbound or loosely bound secondary antibodies were washed out using PBS buffer (pH 7.5), and TMBZ (10 nM)/H_2_O_2_ (5 nM) solution was added to each well (50 μL/well) for 10 min at room temperature. The enzymatic reactions were stopped by adding 10% H_2_SO_4_ solution (50 μL/well), and the absorbance of the solutions was recorded using a microplate reader (Cytation 5, BioTek Instruments Inc., Ontario, Canada) at 450 nm.

### 2.8 Anti-H5N1 HA (Ab 135382) conjugation with self-assembled CAu nanostructures

Target virus-specific antibodies (anti-H5N1 HA Ab 135382) were bound to self-assembled nanostructures of CAu NPs through electrostatic bonds. 1 mL of positively charged (+12.38 eV) CAu nanostructures were mixed with negatively charged (-2.31 eV) HA Ab 135382 antibody (1 μL, 5 ng/mL) and continuously stirred for 2h at room temperature. The mixture was then centrifuged (4000 rpm for 10 min) and washed 3 times with PBS buffer to remove the unbound components. To check the antibody-nanostructure binding, samples were blocked with 2% BSA (100 μL) for 2 h at room temperature. After centrifugation and washing steps, 1 ng/mL (50 μL) of anti-mouse IgG-horseradish peroxidase (HRP) (Santa Cruz Biotechnology, CA) was added to each well followed by incubation at room temperature for 1 h. Upon adding a total of 100 μL TMBZ substrate solution (10 µg/mL, TMBZ, 10% H_2_O_2_ in 100 mM NaOAc, pH 6.0) to each well for 5–30 min at 25°C. Enzymatic color developed (blue color) at this stage, and the reaction was stopped by adding 10% H_2_SO_4_ (100 vμL), and absorbance was recorded at 450 nm with a reference at 655 nm.

### 2.9 Binding of anti-H5N1 NA with carboxyl capped QDs using ELISA

Carboxyl capped CdTe QDs were bound with antibodies through EDC/NHS chemistry. 1 mL of QDs were placed in a 1.5 mL microfuge tube followed by adding 4 mM of EDC, NHS (10 mM) and 1 μL anti-NA antibodies (5 ng/mL), and the mixture was gently stirred at 10°C. To confirm the binding between QDs and antibodies, ELISA was performed as follows: Centrifuged (1000 rpm, 10 min) and washed to remove unbound antibodies or other reagents, then the conjugated part was blocked with 2% BSA for 1h. Anti-mouse IgG-HRP (Santa Cruz Biotechnol., CA) (50 μL, 1ng/mL) was then added in each well after washing steps and then incubation for 1h at room temperature. Samples were washed after centrifugation three times, and 50 μL TMBZ substrate solution (10 μg/mL, TMBZ, 10% H_2_O_2_ in 100 mM NaOAc, pH 6.0) was added to each well and allowed to stand for 5–30 min at 25°C. At last, the reaction was stopped by adding 50 μL of 10% H_2_SO_4_ and the absorbance was recorded at 450 nm.

### 2.10 Chiro-optical detection of Influenza virus A (H5N1)

Chiral based optical sensing experiments were performed after confirming the binding of anti-H5N1 HA (Ab 135382) with CAu nanostructures and QDs using ELISA method. 100 μL of anti-HA Ab-conjugated CAu nanostructure containing varying concentrations of recombinant influenza A (H5N1) in PBS buffer were added to the CD cuvette. Then, 100 μL of anti-NA Ab-conjugated QDs were added, and the CD response was checked. Other influenza viruses were examined in the same manner as a negative control. The samples were excited at 380 nm, and the exciting and the emission slits were at 5 and 5 nm, respectively. Based on the chiro-optical response at different concentrations of target virus, a statistical curve was constructed. In case of clinical virus culture samples, influenza A (H4N6), 50 μL of anti-HA Ab-conjugated CAu nanostructure containing varying concentrations of recombinant influenza A (H4N6) in chicken blood samples were added to the CD cuvette. Then, 100 μL of anti-NA Ab-conjugated QDs were added, and the CD response was recorded. Similar sensing method was followed in case of fowl adenoviruses-9 (FAdVs-9) strain and Infectious bronchitis virus (IBV) detection.

### 2.11 Validation with commercial ELISA kit

A comparison experiment was performed with the commercial avian influenza A H5N1 ELISA kit (Cat. No: MBS9324259, MyBioSource, Inc., San Diego, USA) to validate our proposed method. Different concentrations of virus solution were prepared using sample diluent that was received with the commercial ELISA kit strictly maintaining the manufacturer’s protocol during bioassay. Color developed with different absorbance intensity related to viral concentration in the 96-well plates was recorded at 450 nm. Influenza A (H4N6) virus detection was also validated with commercially available ELISA kit (Cat. No: NS-E10156, Novatein Biosciences, Woburn, USA). The assay procedure was strictly followed as mentioned in the protocol book.

### 2.12 Spectroscopy and structural characterization

Transmission electron microscopy (TEM) images were captured using Tecnai TEM (FEI Tecnai G2 F20, Ontario, Canada). Zeta potential was measured with Zetasizer Nano ZS (Malvern Instruments Ltd., Worcestershire, UK). Circular Dichroism spectrum was recorded using JASCO CD Spectrometer (Model: J-815, Easton, USA).

## 3. Results & Discussions

Artificial chiral plasmonic nanomaterials demonstrate extraordinary rotational ability of polarizing light by engineering intra-particle couplings with light-emitting nanomaterials at nanoscale gap in comparison to naturally occurring chiral materials. This concept was utilized in our study to develop chiro-immunosensor that employs exciton-plasmon interaction in chiral gold nanoparticles (CAu NPs) – CdTe QDs nanohybrids. Two different anti influenza virus surface protein antibodies named anti-HA and anti-NA were bound to CAu NPs and QDs, respectively. Then, a nano-sandwich structure was formed with the addition of target influenza virus A/ Vietnam 1203/04/2009 (H3N2). The amount of CAu NPs and QDs taking part in the nano-assembling process totally depends on the viral concentration. Hence, chiro-optical changes would be influenced by the CAu NPs-QDs nanohybrids. With our proposed sensing concept, it is reasonable to move to a chiro-plasmonic rough surface that has more light scattering properties as well as confined energy in space and enables the creation of a strong coupling regime and increased optical non-linearities through light-matter interactions with QDs.^1,34^ Such strong localization of energy in nanoscale gap may provide specific spatial positions to tune and further enhance the chiral property of CAu NPs efficiently.

In order to fully exploit the nanostructured morphology, a series of differently shaped (hetero-structured) CAu NPs were prepared by varying L(+) ascorbic acid concentration during the reaction process. As shown in Figure **1**, different sizes and morphologies of CAu NPs were obtained by using various concentrations of L(+) ascorbic acid in 20 mM HAuCl_4_ solution. Prolate shaped CAu NPs were obtained with a diameter of 50 nm at 0.1 M L(+) ascorbic acid (Fig. 1A). As the concentration of L(+) ascorbic acid shifted from 0.1 M to 0.005 M (Figure 1 A–D), the CAu NPs structure ranged from well-defined simple flower-like (Fig. 1B), urchin (Figure 1C), and dendritic-type morphologies (Fig.1D). It should be noted that urchin shaped CAu NPs with many spikes were obtained when the concentration of L(+) ascorbic acid was 0.01M (Fig. 1C). Unfortunately, though dendritic-shaped CAu NPs contained many arms and branches on their structural morphology, spectroscopic study showed a very weak plasmonic peak (Fig.2). The plasmonic peaks obtained from prolate, flower, and urchin shaped CAu NPs were located at 548 nm, 565 nm, and 590 nm, respectively. Among them, urchin shaped CAu NPs was chosen for further experiments due to the numerous spikes on their surface as well as a broadened plasmonic peak, which offers the possibility of having maximum overlap, and coupling with the excitonic wavelength of QDs, which may strengthen the optical activity.

**Figure 1:**
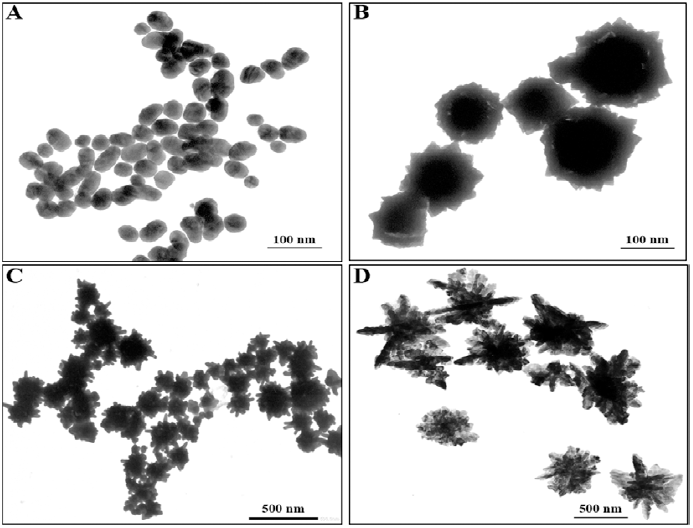
Transmission electron microscopic image of the developed CAu NPs; (A) Prolate-like, (B) Flower-like, (C) Urchin-like & (D) Dendritic shaped CAu NPs.

**Figure 2:**
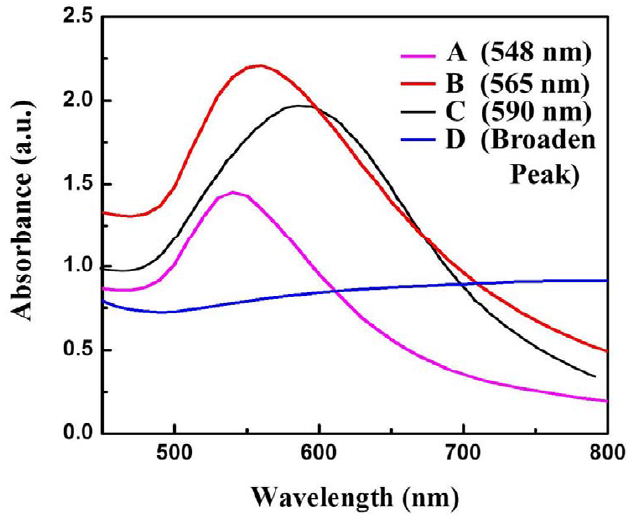
UV-visible spectra of different shaped CAu NPs; (A) Prolate-like, (B) Flower-like,(C) Urchin-like & (D) Dendritic shaped CAu NPs.

Enhanced localized surface plasmon resonance (LSPR) could potentially serve as an effective platform to prove that the hetero-structured nanoparticles have enough sensitivity for bioanalytical detection.^35^ With this concept in mind, the self-assembled structure of star-shaped nanoparticles was prepared by the addition of CTAB that served as a ‘glue’ to link the {100}facets of two adjacent CAu NPs, which in-turn aided in organizing the chiral gold nanostructure into a short chain.^36^ Star-shaped CAu nanostructures with varying morphology is shown in Figure 4, with spike length and diameter of 10 nm and 2 nm, respectively (Fig. 3F).

**Figure 3:**
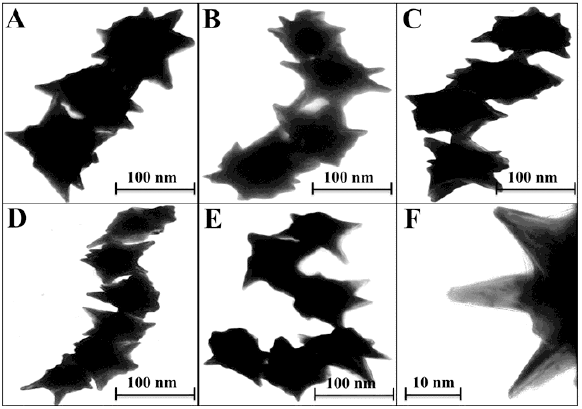
Transmission electron microscopic image of the self-assembled chiro-plasmonic nanostructures; (A) short straight chain, (B) left turned chain, (C) right turned chain, (D) long chain with short angled curve chain, (E) long chain with high angled curve chain & (F) close-up image of spikes of CAu NPs.

**Figure 4:**
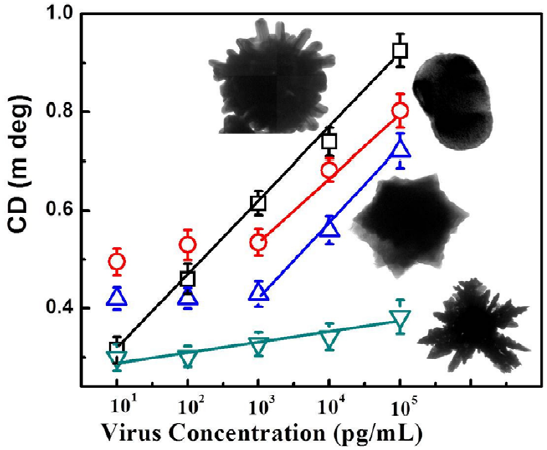
Comparison of the detection performance of different shaped CAu NPs; CD spectrum Vs target virus sensing results.

CdTe NPs was chosen to make nanohybrids with CAu NPs due to its strong emission properties and energy overlap possibility between the excitonic and plasmonic states of the resulting nanohybrids. Photoluminescence spectra (PL) and ultra-violet (UV) spectra of QDs are shown in Fig. S1A; the emission peak was located at 710 nm. Particle size and concentration were 6.5 nm and 3.7× 10^−7^ M, respectively, calculated based on Peng equation.^37^

The specificity of anti-influenza A (H5N1) virus hemagglutinin (HA) antibody Ab 135382 against recombinant influenza virus A (Avian/Vietnam/1203/04) (H5N1) was confirmed using a conventional ELISA method. Figure S1B shows that the optical density obtained due to enzymatic activity for the target virus/Ab 135382 HA/HRP-conjugated secondary antibody/ TMBZ–H_2_O_2_ complex, and the target virus/anti-H5N1 NA/HRP-conjugated secondary antibody/ TMBZ–H_2_O_2_ complex was higher than those of anti-H5N2 HA and anti-H7N9 HA Ab, implying the specificity of Ab 135382 towards recombinant influenza virus A (Avian/Vietnam/1203/04) (H5N1). The higher optical density obtained for anti-H5N1 Ab 135382 and CAu nanostructures complex in ELISA results confirmed the successful binding between anti-H5N1 Ab 135382 and CAu nanostructures (Fig. S2A).

The binding between anti-H5N1 Ab 135382 and CdTe QDs was confirmed by FTIR spectrum. As shown in Fig. S2B, FTIR bands found at 3500–3700 cm^−1^ for amide N–H stretching represents the chemical binding between Ab 135382 and CdTe QDs, whereas only carboxylic acid O-H stretching bands appeared for CdTe QDs alone. The influence of the structural morphology on the sensitivity of a chiro-sensor was further investigated to enable the selection of the optimized shape of the CAu NPs for the bioassay platforms. Four typical CAu NPs structures, prolate shaped CAu NPs, flower-shaped CAu NPs, spiky-like CAu NPs and dendritic shaped CAu NPs were selected to construct nanohybrids with QDs through immune-reaction for a chiral biosensor for avian influenza virus detection. It was found that the optical response of the biosensors was largely dependent on the total amount of analytes assembled on these CAu NPs. As shown in Figure 4, the linear detection ranges of CAu NPs were drastically influenced by the nanostructure’s shape. The detection range of urchin-like CAu NPs was 10 pg to10 μg/mL, whereas for prolate and flower shaped CAu NPs, the range was between 1 ng to 10 μg/mL.

Experimental results revealed that chiro-optical response from dendritic shaped CAu NPs was not significant; probably due to a weak plasmonic peak. In general, rough nanostructures have higher analytes capture efficiency in the bioassay applications than that of the smooth nanostructures, and this ultimately influences the detection limit of a biosensor. Compared with other CAu NPs, urchin shaped nanostructures possessed higher surface to volume ratio as well as stronger plasmonic peak; therefore, we chose urchin shaped CAu NPs to construct the self-assembled chiro-plasmonic nanostructures.

Nano-sandwich assembly was formed via the addition of various concentrations of recombinant influenza virus A (H5N1) to antibody conjugated self-assembled CAu nanostructure and QDs solutions. Chiro-optical response and plasmonic response of viral concentration were carefully examined (Fig. 5A & 5B), and a calibration curve for chiro-optical detection of avian influenza virus was constructed. The sensitivity of the bioassay was found up to 1 pg/mL (Fig. 5C), 10 times more sensitive than the non-assembled urchin-like CAu NPs. Most probably, multi-times scattering of incident light on the plasmonic rough surface confined lots of energy in space, which ultimately influence the chirality of CAu nanostructures. A plasmon-based bioassay was performed using the same standard viral concentrations, but the plasmonic response did not significantly correlated with the viral concentrations. In the case of conventional ELISA method, the sensitivity was found to be 1 ng/mL level.

**Figure 5:**
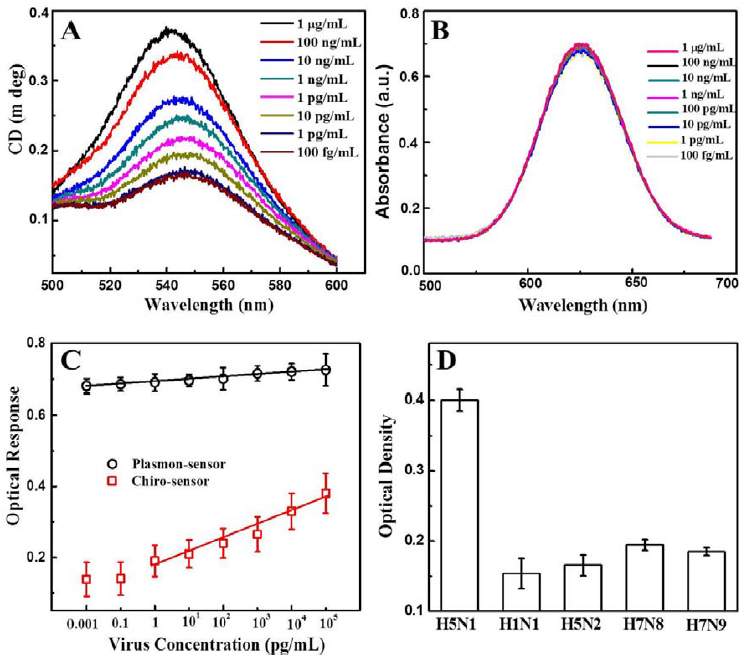
Detection of the avian influenza virus; (A) Chiro-optical spectra of different concentration virus detection, (B) Plasmonic spectra of different concentration virus detection, (C) Analytical calibration curves relating the intensity of circular dichroism (CD) and plasmonic spectra & (D) Selectivity of proposed bioassay.

The selectivity test of the proposed bioassay was implemented with other virus strains, namely H1N1, H5N2, H7N8 and H7N9; a significant chiro-optical response (8~9-fold higher) was obtained with the target avian influenza virus A (H5N1) in comparison to the others, revealing that the current bioassay is sufficiently selective for the detection of target avian influenza virus A (H5N1) (Fig. 5D).The sensitivity of the developed chiro-optical sensor was validated with the commercial colorimetric detection kit for avian influenza A (H5N1) (Table 1 & Fig. S3). The visual color response of the commercial kit was up to 1 ng/mL, indicating that the chiro-optical response-based bioassay is more sensitive than the commercial kit.

**Table 1:**
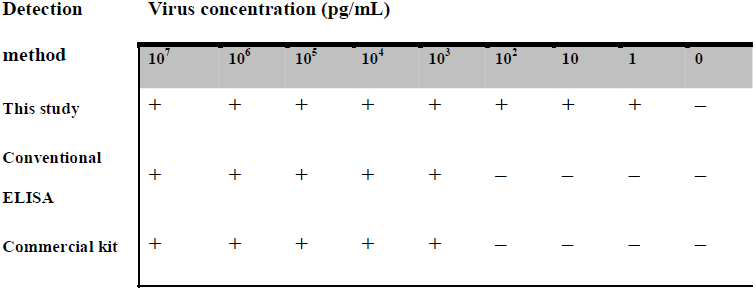
Comparison of avian influenza virus A (Avian/Vietnam/1203/04) (H5N1) detection using different methods

We also explored the practical application of the developed chirosensor in the PBS buffer and in chicken blood samples for influenza A (H4N6) virus detection that represents a complex biological mixture. The detection of the Avian influenza A (H4N6) virus concentration is dependent on the amount of CAu nanostructures and QDs presents in nano-assembled structure and hence is directly associated to the optical activity of the resulting dispersions. Upon confirming the anti-H4N6 antibody specificity towards the real avian influenza A (H4N6) virus (Fig. S4) and its binding confirmation with CAu nanostructure and QDs (Fig. S5); a calibration curve for chiroplasmonic detection of influenza A (H4N6) virus was obtained by the standard dilution method (Fig. 6A). The linear range of the influenza A (H4N6) virus detection in PBS buffer was found to be from 100 to 0.01 HAU/50 μL with the limit of detection 0.0268 HAU/50 μL. The proposed chirosensor exhibited its detection ability at complex media with the linear range of 100 to 0.01 HAU/50 μL; the limit of detection was 0.0315 HAU/50 μL calculated by standard deviation method. Here, bovine serum albumin (BSA) and influenza virus A (H5N1) were used a control. The validated ELISA results confirmed the superiority of the proposed technique in terms of sensitivity over the commercial kit (Table S1). FAdVs-9 viruses in blood samples were detected followed by confirming the successful binding of anti-FAdVs antibody through ELISA method (Fig. S6). The sensitivity of the chiral response based bioassay for FAdVs-9 detection showed up to 50 PFU/mL with the limit of detection of 33.64 PFU/mL (Fig. 6B) and the assay confirmed the highly selectivity as well (Fig. S7).

**Figure 6:**
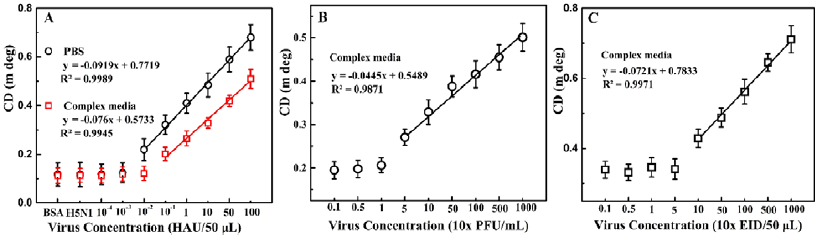
Detection of the multiple virus culture samples; (A) avian influenza virus A (H4N6); Closed Circles (black) and square (red) represent the sensing results of A (H4N6) detection in PBS buffer and complex matrix (chicken blood) respectively, (B) fowl adenovirus (FADV) and (C) Infectious bronchitis virus (IBV) detection in chicken blood media.

Our proposed bioassay system was further applied for infectious bronchitis virus (IBV) detection in blood samples after confirming successful binding of CAu nanostructures and QDS with IBV specific antibody (Fig. S8). The chiroptical response of IBV detection was observed to be in the range of 10^2^~10^4^ EID/50 μL with the limit of detection of 47.91 EID/50 μL (Fig. 6C) and the assay confirmed selectivity for only the target analytes (S9).Thus, the proposed chiral-based biosensor enable the utilization of this technique in real life applications.

## 4. Conclusions

Chiral biosensors will be at the forefront of the bionanotechnology field in the near future due to the ultra-sensitivity and rapid response time. Our work demonstrates a framework and a method to achieve enhanced sensitivity using chiral metal nanoparticles–quantum dot nanocomposites for detecting bioanalytes in clinical samples. While the proposed sensing method was applied to the colloidal CAu NPs and self-assembled CAu nanostructure; the latter showed better detection performance due to the multipole plasmonic scattering effects on the rough CAu nanostructure. Sensitivity on the order of pico-gram level was achieved, which is superior than the conventional ELISA methods, the plasmonic biosensor and the commercial kits. The sensing performance of the proposed method was also retained in complex biological media for avian influenza A (H4N6) virus, fowl adenovirus and coronavirus. The simplicity and more importantly, the concept of optical rotation and energy coupling phenomena in the chiral plasmon-exciton systems may have academic significance, opening new doors for chiral bionanosensor development and further studies of chiro-optical memory, chiral catalysis, and light-emitting diodes.

## Acknowledgements

The authors sincerely thank the Natural Sciences and Engineering Research Council of Canada (400705) for funding this study. The authors thank Professor Rod Merrill of the Molecular and Cell Biology department of the University of Guelph for providing access to the circular dichroism equipment.

## Notes and references

1. M. Achermann, J. Phys. Chem. Lett., 2010, 1, 2837–2843.

2. E. Cohen-Hoshen, G. W. Bryant, I. Pinkas, J. Sperling and I. Bar-Joseph, Nano Lett., 2012, 12, 4260–426.

3. J. Lee, P. Hernandez, J. Lee, A. O. Govorov and N. A. Kotov, Nat. Mater., 2007, 6, 291–295.

4. F. Todisco, S. D'Agostino, M. Esposito, A. I. Fernández-Domínguez, M. D. Giorgi, D. Ballarini, L. Dominici, I. Tarantini, M. Cuscuná, F. D. Sala, G. Gigli and D. Sanvitto, ACS Nano, 2015, 9, 9691–9699.

5. R, Jiang, B. Li, C. Fang and J. Wang, Adv. Mater., 2014, 26, 5274–5309.

6. Q. Zhang, L. Lee, J. B. Joo, F. Zaera, and Y. Yin, Acc. Chem. Res., 2013, 46, 1816–1824.

7. V. K. Valev, J. J. Baumberg, C. Sibilia and T. Verbies, Adv. Mater., 2013, 25, 2517–2534. 462

8. M. H. Alizadeh and B. M. Reinhard, ACS Photonics, 2015, 2, 942–949.

9. Y. Zhao, A. A. E. Saleh and J. A. Dionne, ACS Photonics, 2016, 3, 304–309.

10. J. Butet, P. Brevet and O. J. F. Martin, ACS Nano, 2015, 9, 10545–10562.

11. M. Scha□ferling, X. Yin, N. Engheta and H. Giessen, ACS Photonics, 2014, 1, 530–537.

12. T. Hu, B. P. Isaacoff, J. H. Bahng, C. Hao, Y. Zhou, J. Zhu, X. Li, Z. Wang, S. Liu, C. Xu, J. S. Biteen and N. A. Kotov, Nano Lett., 2014, 14, 6799–6810.

13. W. Ma, H. Kuang, L. Wang, L. Xu, W. Chang, H. Zhang, M. Sun, Y. Zhu, Y. Zhao, L. Liu, C. Xu, S. Link and N. A. Kotov, Sci. Rep., 2013, 3, 1934.

14. XWu, L. Xu, L. Liu, W. Ma, H. Yin, H. Kuang, L. Wang, C. Xu and N. A. Kotov, J. Am. Chem. Soc., 2013, 135, 18629–18636.

15. Y Zhao, L. Xu, W. Ma, L. Wang, H. Kuang, C. Xu and N. A. Kotov, Nano Lett., 2014, 14, 3908–3913.

16. X. Shen, A. Asenjo-Garcia, Q. Liu, Q. Jiang, F. J. G. D Abajo, N. Liu and B. Ding, Nano Lett., 2013, 13, 2128–2133.

17. X. Lan, X. Lu, C. Shen, Y. Ke, W. Ni and Q. Wang, J. Am. Chem. Soc., 2015, 137, 457–462.

18. J. Cheng, G. L. Saux, J. Gao, T. Buffeteau, Y. Battie, P. Barois, V. Ponsinet, M. Delville, O. Ersen, E. Pouget and Reiko Oda, ACS Nano, 2017, 11, 3806–3818.

19. W. Chen, A. Bian, A. Agarwal, L. Liu, H. Shen, L. Wang, C. Xu and N. A. Kotov, Nano Lett., 2009, 9, 2153–2159.

20. A. J. Mastroianni, S. A. Claridge and A. P. Alivisatos, J. Am. Chem. Soc., 2009, 131, 8455–8459.

21. A. Guerrero-Martínez, B. Auguié, J. L. Alonso-Gómez, Z. Džolić, S. Gómez-Graña, M. žinić, M. M. Cid, L. M. Liz-Marzán, Angew. Chem., 2011, 123, 5613–5617.

22. Z. Fan and A. O. Govorov, Nano Lett., 2010, 10, 2580-2587.

23. J. Zho, S. Tang, J. Storhoff, S. Marla, Y. P. Bao, X. Wang, E. Y. Wong, V. Ragupathy, Z. Ye and I. K Hewlett, BMC Biotechnol., 2010, 10, 74.

24. J. Lum, R. Wang, K. Lassiter, B. Srinivasan, D. Abi-Ghanem, L. Berghman, B. Hargis, S. Tung, H. Lu and Y. Li, Biosens. Bioelectron., 2012, 38, 67–73.

25. S. W. Thor, H. Nguyen, A. Balish, A. N. Hoang, K. M. Gustin, P. T. Nhung, J. Jones, N. N. Thu, W. Davis, T. N. T. Ngoc, Y. Jang, K. Sleeman, J. Villanueva, J. Kile, L. V. Gubareva, S. Lindstrom, T. M. Tumpey, C. T. Davis and N. T. Long, PLOS ONE, 2015, 10, e0133867.

26. C. Li, D. Cao, Y. Kang, Y. Lin, R. Cui, D. Pang and H. Tang, Anal. Chem., 2016, 88, 4432–4439.

27. L. Deng, S. Sharif and É. Nagy, Clin. Vaccine Immunol., 2013, 20, 1189–1196.

28. H. Grgić, D. B. Hunter, P. Hunton and É. Nagy, Can. J. Vet. Res., 2008, 72, 403–410.

29. M. St Paul, N. Barjesteh, J. T. Brisbin, A. Villaneueva, L. R. Read, D. Hodgins, É, Nagy and S. Sharif, Viral Immunol. 2014, 27, 167–173.

30. H. S. Alexander, P. Huber, J. Cao, P. J. Krell and É Nagy, J. Virol. Methods, 1998, 74, 9–14.

31. H. Grgić, D. B. Hunter, P. Hunton and É. Nagy, Can. J. Vet. Res., 2008, 72, 403–410.

32. J. J. Gelb, 4th ed. The American Association of Avian Pathologists, Dubuque, Iowa: Kendall/Hunt Publ, 1998, 169–175.

33. L. J. Reed and H. Muench, Am. J. Hyg. 1938, 27, 493–497.

34. B. Xiao, S. K. Pradhan, K. C. Santiago, G. N. Rutherford and A. K. Pradhan, Sci. Rep., 2016, 6, 24385.

35. H. Yockell-Lelievre, F. Lussier and J. Masson, J. Phys. Chem. C, 2015, 119, 28577-28585.

36. Y. Yang, S. Matsubara, M. Nogami, J. Shi and W. Huang, Nanotechnology, 2006, 17, 2821–2827.

37. W. W. Yu, L. Qu, W. Guo and X. Peng, Chem. Mater., 2003, 15, 2854–2860.

